# Prediction of acoustic tinnitus suppression using resting state EEG: An explainable AI approach

**DOI:** 10.1101/2024.04.16.589690

**Authors:** Payam S. Shabestari, Stefan Schoisswohl, Zino Wellauer, Adrian Naas, Tobias Kleinjung, Martin Schecklmann, Berthold Langguth, Patrick Neff

## Abstract

Tinnitus, characterized by the perception of sound without an external source, affects a significant portion of the population and can lead to considerable individual suffering, yet understanding of its suppression remains limited. Understanding neural traits of tinnitus suppression may be crucial for developing accurate predictive models in tinnitus research and treatment. This study aims to classify individuals capable of brief acoustic tinnitus suppression (BATS; also known as residual inhibition) based on their independent resting state EEG (n=102), exploring the classification’s robustness on various sample splits, and the relevance of resulting specific EEG features in the spirit of explainable AI. A comprehensive set of EEG features, including band power in standard frequency bands, spectral entropy, aperiodic slope and offset of the power spectrum, and connectivity, was included in both sensor and source space. Binary classification of the BATS status was performed using a comprehensive set of standard classifiers and Pearson correlation for feature selection, which addresses multicollinearity, avoiding complex dimensionality reduction techniques. Feature importance was assessed using Gini impurity metrics, allowing interpretation of the directionality of identified neural features. The Random Forest model showed the most consistent performance, with its majority voting mechanism effectively reducing overfitting and providing reliable predictions, and was therefore chosen for subsequent feature interpretation analysis. Our classification task demonstrated high accuracy across the various BATS split thresholds, suggesting that the choice of threshold does not significantly influence the underlying pattern in the data. We achieved classification accuracies of 98% for sensor and source models and 86% for the connectivity model in the main split. Looking at identified important features, our findings align with and extend existing neuroscience research in tinnitus by discovering highly specific and novel neural features in naive resting-state data predictive of BATS. Gamma power is identified as the most important feature in the sensor model, followed by alpha power, which fits current models of sensory processing, prediction, and updating (gamma) as well as inhibitory (alpha) frameworks. The overall spectral shape of the EEG power spectrum tends to be more normal in +BATS individuals, as reflected in the aperiodic offset and slope features. In the source model, important features are lateralized in that the gamma feature is more prominent in the left core auditory network, whereas the alpha feature is distributed more sparsely over the right hemisphere in line with auditory attention data. Furthermore, we identified several hotspots in the temporal, insular, parietal, parahippocampal, medial prefrontal, and (posterior) cingulate cortex implicated in sensory processing, gating, attention, and memory processes. Relevant network features were found in a hyperconnected bilateral auditory network (within the network), while the full auditory network was hyperconnected to limbic regions (between networks), which may reflect an intact sensory gating mechanism aiding tinnitus suppression. This study’s implications extend to improving the understanding and prediction of tinnitus loudness perception and tinnitus distress as well as its (acoustic) suppression. Furthermore, our approach underscores the importance of careful feature selection, model choice, and validation strategies in analyzing complex neurophysiological data.

## Introduction

Chronic subjective tinnitus is defined as the persistent and conscious auditory perception of tonal or composite noise in the absence of an equivalent external physical acoustic source, which can evolve into a more complex syndrome termed ‘tinnitus disorder’ marked by high levels of tinnitus-related distress [1, 2]. With a prevalence of about 14.4%, tinnitus is a common condition in the global population with many of those affected experiencing severe burden [3], and suffer from several comorbidities such as depression or anxiety disorders [4]. Currently, no effective treatment for tinnitus is established or on the horizon and the available treatment approaches only focus on secondary symptoms such as quality of life management [5–7]. Typically, tinnitus is thought to evolve as a consequence of noise trauma and/or hearing loss [8], whereby the resulting lack of peripheral auditory input provokes maladaptive pathological changes in the auditory pathway as well as the central nervous system putatively responsible for the perception of the auditory phantom sound perception tinnitus [9–11]. These pathological alterations further translate into distinctive tinnitus-related spontaneous brain activity patterns; here, increased activity in the delta and gamma frequency bands and decreased activity in the alpha frequency band in (sensory) auditory cortical regions have been reported by electroencephalography (EEG) or magnetoencephalography (MEG) by several research groups [12–15]. Moreover, tinnitus-related alterations in functional global and modality-specific networks have been reported [16]. Generally, these alterations include increased connectivity within and between the auditory network, the default mode network, the attention networks, and the visual network.

In past studies, various machine learning approaches were applied to differentiate the tinnitus population from healthy controls using resting state EEG data with high accuracy [17–20]. However, to this day, machine learning studies concerning active manipulation of tinnitus have not been carried-out within the tinnitus population.

A large portion of individuals with tinnitus (60-80%) are capable of undergoing temporary suppression of the subjective tinnitus perception to some degree following sound stimulation with either white noise, sine tones, or various (complex) modulated or filtered stimuli [21–27]. Brief Acoustic Tinnitus Suppression (BATS), established as “residual inhibition” or “forward masking” [28, 29], is theorized to result from a temporary recovery of imbalanced inhibitory and excitatory neuronal activity in the auditory cortex and/or reduced firing of neurons along the auditory pathway [30]. While studying the subcortical auditory pathway below the brainstem in human participants is challenging, so far only three studies focused on cortical activity related to BATS on a group level, besides three single case studies showing heterogeneous findings [31–33]: Kahlbrock and Weisz [34] observed a decline in pathologically enhanced activity in the delta frequency range, whereas King and colleagues’s study indicated elevated power spectral density concerning gamma and alpha activity [35]. Similarly, we could demonstrate in our former study [36] that participants who experience BATS had enhanced alpha activity in general compared to participants without BATS. In contrast to King and colleagues’ study, however, we observed reduced gamma band amplitudes. This observation further emphasizes specific oscillatory signatures of tinnitus patient subtypes related to the ability to induce short-term tinnitus suppression via acoustic stimulation. Currently, there seems to be convergence regarding the role of alpha in BATS with most studies reporting an increase while in other frequency bands, especially gamma, the results are diverging and partly contradicting.

To the best of our knowledge, no former study attempted to predict tinnitus suppression, ‘off-states’, or specifically BATS from (naive) resting state M/EEG data. In contrast to former studies in BATS probing short-term state-like neural responses during BATS or classification thereof, this approach may elucidate how individual trait-like or phenotypic neural signatures influence the ability to suppress tinnitus. We therefore consider elaboration on trait-specific (oscillatory) brain activity patterns associated with BATS to be of high interest, given the potential to accurately identify individuals with the ability to acoustically suppress their tinnitus. Automatic, high-accuracy classification and insights gained from resulting, distinctive features would enable us to better understand the BATS phenomenon and basic mechanism of tinnitus on the neural level. Explainable AI has the potential to surpass traditional analysis methods by enabling a comprehensive examination of models and important features [37]. Furthermore, this approach could foster (objective) diagnostic options, tinnitus subtyping, and identification of individual treatment options like sound therapies [38]. Related to that, it was recently shown that certain EEG features such as frequency band power and functional connectivity could predict treatment response to a sound-based intervention with 98-100% accuracy [39].

Currently, it remains unclear which trait-like factors or signatures of (oscillatory) brain activity and connectivity can predict BATS. Hence, the objective of the present work is to apply automatic classification algorithms to evaluate if distinctive EEG sensor, source, and connectivity features are predictive of BATS.

## Materials and methods

### Data sets

The EEG and behavioral data used in this study were sourced from two distinct labs at the universities of Regensburg, Germany, and Zurich, Switzerland. The Regensburg dataset encompassed 79 participants who actively participated in two neurobehavioral experiments investigating BATS with EEG. These experiments received ethical approval from the internal ethics review board of the Faculty of Medicine, Regensburg, under reference numbers 17-819-101 and 18-1054-101. The Zurich validation set consisted of 29 participants partaking in a neuromodulation study where EEG and BATS were assessed during baseline measurements. Ethical approval for the Zurich study was obtained from the Cantonal Ethics Committee (KEK, Zurich; BASEC-Nr. 2020-02027). All individuals included in the studies pertaining to the dataset at hand provided informed consent for both their participation in the studies and the utilization of their data for future analyses. The experiments were conducted in strict compliance with the ethical principles outlined in the Declaration of Helsinki.

For more comprehensive details and descriptive statistics related to the data sets, readers are referred to the supplementary material tables S1 Table and S2 Table.

### Feature extraction

An overview of our method pipeline is shown in Fig 1. Following the preprocessing and epoching of the EEG data, a set of frequency domain features, comprising oscillatory power estimation, non-oscillatory parameters, and information measures, was extracted for each EEG epoch, with calculations performed individually at the level of each electrode and brain region. The features computed per electrode consisted of: Sensor space spectral power values averaged within the five canonical M/EEG frequency bands (comprising 310 features), average spectral Shannon entropy (comprising 310 features) [40, 41], and non-oscillatory parameters, such as the slope and offset of power spectral density at each electrode (comprising 124 features) [42]. For each designated brain region extracted from Desikan-Killiany [43] atlas (source space), the features computed encompass the average spectral power across five distinct frequency bands within the epochs (comprising 340 features). See the supplementary section for a detailed explanation of how each feature set was computed. The (standard) frequency bands utilized for computing features encompass: Delta (0.5-4.5 Hz), Theta (4.5-8.5 Hz), Alpha (8.5-13.5 Hz), Beta (15-30 Hz) and Gamma (30-80 Hz) [44].

**Fig 1.**
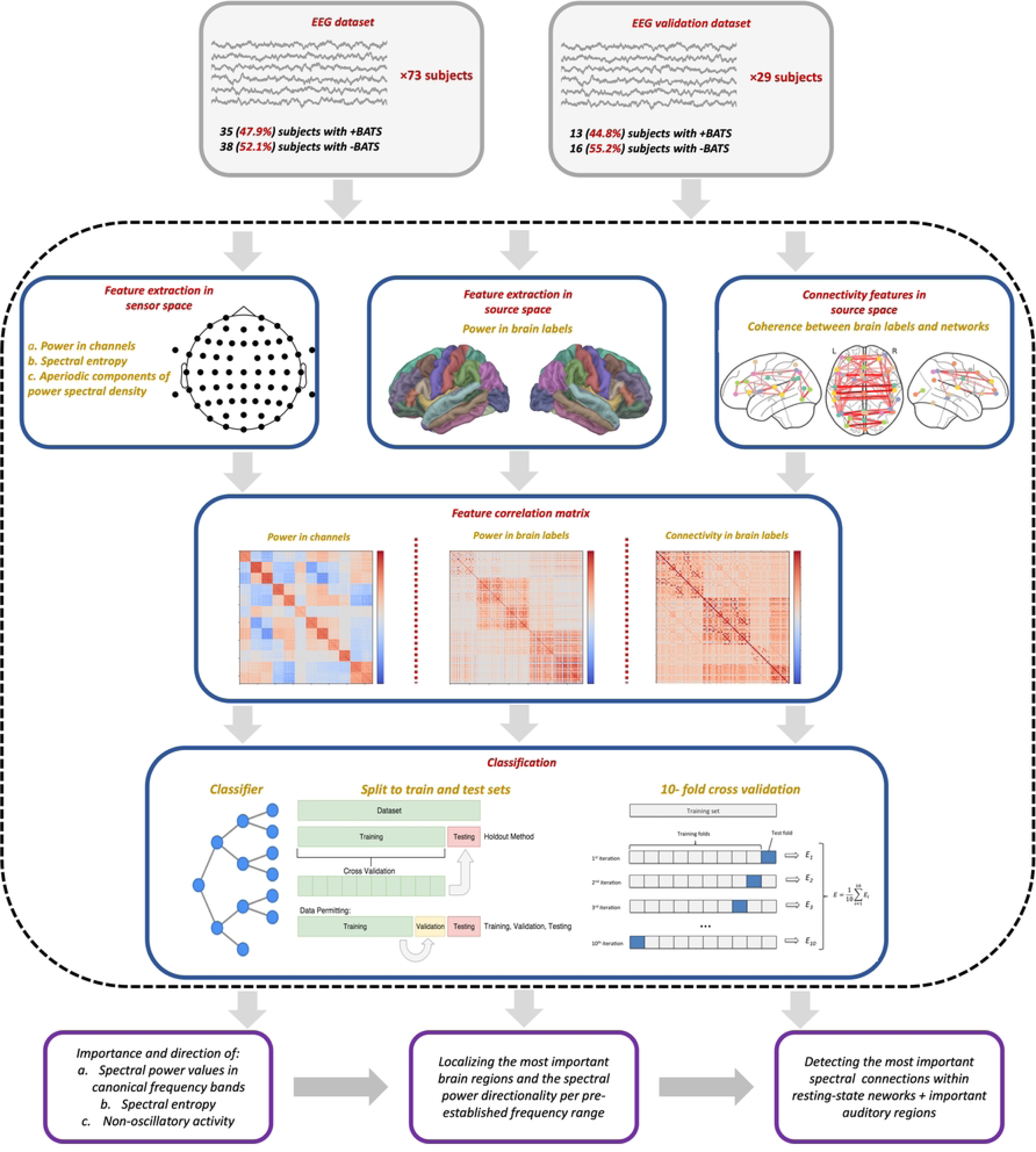
Overview of the analysis pipeline. EEG data collected from 73 participants (main dataset) was divided into 2-second epochs after preprocessing. Sensor space features with high correlation were merged using Pearson correlation. Data was partitioned into training and test sets, and a 10-fold cross-validation was performed on the training set. Further analysis involved ranking features based on their importance and exploring their directional impact. A parallel process was applied to the epochs using features computed in the source space. The outcomes of these analyses were utilized to calculate spectral connectivity features in each frequency band and between brain regions (labels) in the source space. Furthermore, all procedures were performed on an additional, independent EEG dataset of 29 participants to validate and benchmark the results.

Due to the intrinsic correlation between spectral power values in the sensor space and source space, we divided the features into two distinct sets. Feature set 1, which pertains to the sensor space, was employed to explore the significance and direction of spectral power and spectral entropy in canonical frequency bands, as well as non-oscillatory activity in predicting BATS. By exploiting the classification results of feature set 1, it was narrowed-down into feature set 2 in source space, focusing solely on spectral power values associated with brain regions. This procedure will help to investigate the specific contribution of individual brain regions (or labels) in the prediction of BATS.

Further, coherence metrics across frequency bands and brain regions identified as important through source space analysis were computed (refer to Section S8 Text in the supplementary materials for more details). For the analysis of functional connectivity patterns across different resting-state brain networks, the Desikan-Killiany atlas was utilized [43] and brain regions were organized into networks as delineated in the supplementary Table S3 Table. Seven large-scale functionally segregated networks [45, 46] were categorized, including visual (VSN), somatomotor (SMN), dorsal attention (DAN), ventral attention (VAN), limbic (LBN), frontoparietal (FPN), and default mode (DMN) networks. Recognizing the particular importance of the auditory network (AUN) in tinnitus, we incorporated nine distinct sub-networks along with most contributing brain regions detected within the AUN network. Labels for the left or right hemisphere were included if the corresponding label in the other hemisphere was missing to account for the brain’s bi-hemispheric functional organization.

### Classification pipeline

Features with correlation coefficients exceeding 0.9 were merged using Pearson correlation. The selected set of features significantly improved the model’s interpretability and its ability to generalize across different datasets. Then, the classification process was performed by shuffling the epochs and then randomly splitting them into a 70% training and a 30% test set. Following this initial split, a 10-fold cross-validation procedure was carried-out exclusively on the training set, and finally, the classification algorithm’s accuracy was computed using the test set as follows:

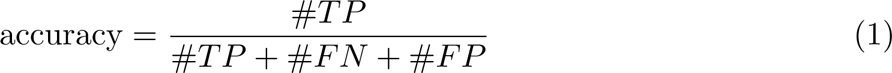

where #*TP* represents the count of true positives, signifying correctly classified epochs from individuals with BATS. #*FN* corresponds to the count of false negatives, encompassing misclassified epochs from individuals with +BATS, and #*FP* denotes the count of false positives, including individuals incapable of persistently suppressing tinnitus (-BATS) but have been incorrectly assigned to the other class. In our analysis, we employed 10 widely recognized classifiers from the scikit-learn python package [47]. These classifiers comprise: Random Forest (RF) [48], Gradient Boosting [49], Quadratic Discriminant Analysis (QDA) [50], Naive Bayes [51], Decision Tree [52], Radial Basis Function (RBF) kernel SVM [53], Gaussian Process [54], k-nearest neighbors [55], Convolutional Neural Network [56] and linear SVM [57]. By utilizing such a diverse set of classifiers, it was evaluated if the classification task is robust across classifiers and the different thresholds for BATS.

After benchmarking the set of classifiers, the best performing model was selected and subjugated to feature importance analysis. Feature importance was determined based on the mean and standard deviation of the reduction in impurity within each tree, known as Gini impurity metric (see section S8 Text in supplementary materials). Subsequently, we retrained the model, using only the top 100 features identified through this process. To assess the significance and directional impact of features (sensor space, source space, or spectral connections) in predicting the two classes (i.e., +BATS and -BATS), we utilized the SHAP (SHapley Additive exPlanations) Python package [58].

### Validation

To validate and benchmark our results, we utilized a distinct validation dataset, which is detailed in the Data sets section. The same methodology as detailed in sections Feature extraction and Classification pipeline was applied to the validation dataset, including the computation of predefined feature sets both in the sensor and source space as well as spectral connectivity measures. For classification, a BATS threshold of −1 was employed to categorize participants into two groups: those who did show acoustical suppression of their tinnitus (with values less than or equal to −1) and those who did not (with values larger than −1). Note, the scale ranged from 0 indicating no suppression to −5 indicating full suppression. Moreover, we computed models where the BATS labels (i.e., +BATS and -BATS) of the data split were randomly shuffled so each label consisted of a mixture of true and false labels (50% mixture). This allowed for validation of our models and the related ground truth assumption, namely, the ability to suppress tinnitus based on individuals’ self-reports.

## Results

In the analysis of the main dataset, statistically significant higher Minimum Masking Levels (MML) were observed in the -BATS group compared to the +BATS group. This result indicates a potential correlation between the ability to acoustically suppress tinnitus and the MML (mean difference = 7.63 dB, S2 Table). Additionally, while not reaching the level of statistical significance, there was a trend towards higher tinnitus loudness levels in the -BATS group (mean difference = 8.58 dB), which is in line with the MML finding and suggests a relationship between tinnitus maskability during sound presentation and residual inhibition after sound presentation.

### Sensor space

As a result of the feature correlation check, 92 features, which accounted for 12.4% of the original 744, were excluded due to their high correlation. We furthermore assessed the accuracy of our models on test data by setting the BATS threshold to five different values (here: perceived tinnitus loudness during +BATS): 90, 80, 70, 60, and 50, as illustrated in Fig 2. For consistency in subsequent classification tasks conducted in the source space and connectivity analysis, we adopted RF as our standard classifier and set the threshold for +BATS to 90 (see for more details). Notably, our ancillary randomly-shuffled label models analysis for this threshold resulted in an accuracy of 51.7% for the RF model, providing clear evidence for the feasibility of our choice of ground truth in this analysis. After selecting the top 100 most important features and retraining the model, we achieved an accuracy of 98.6% for the RF model on the test data. Subsequently, by assessing the importance of these features, we discovered that the power spectrum averaged over the gamma and alpha frequency bands exerted the most significant influence on the model’s predictions for both classes, namely +BATS and -BATS, as depicted in Fig 3.A. Investigating the directional impact of these features, we observed that spectral power values in the gamma frequency range had a bidirectional effect on predicting both classes, with a tendency of high gamma feature values predicting +BATS. Moreover, higher alpha feature values apparently exert a more significant impact on predicting individuals with +BATS, suggesting that individuals with tinnitus who show higher alpha activity during an independent EEG resting-state measurement are more likely to successfully show inhibition of their tinnitus. Channels identified as important contributors by the classification are depicted in Fig 3.B and mostly covering auditory, sensory, and/or attentional networks. Yet, given well-known issues of volume conduction, diffusion, and smearing in M/EEG sensor level localization, source-localized data, as presented in the next section, is more feasible for further interpretation. Entropy values were calculated in order to extend the assessment of power values by contributing measures of orderliness or informational value. Looking at the impact on the model output (SHAP values) in Fig 3.A and D, entropy feature values confirm the bidirectional outcome for gamma and the positive influence of alpha power on tinnitus suppression by showing an inversion of the distribution of the power effects. Furthermore, power values in alpha and gamma are negatively correlated, shown in Fig 3.D in the right-most subplot. Finally, aperiodic parameters complement the results of the feature set showing that +BATS is related to lower aperiodic offsets and steeper slopes (resulting in a larger area under the curve), which reflects more periodic (i.e., oscillatory) activity or ‘normal’ power frequency spectrum in individuals with +BATS (Fig 3.C).

**Fig 2.**
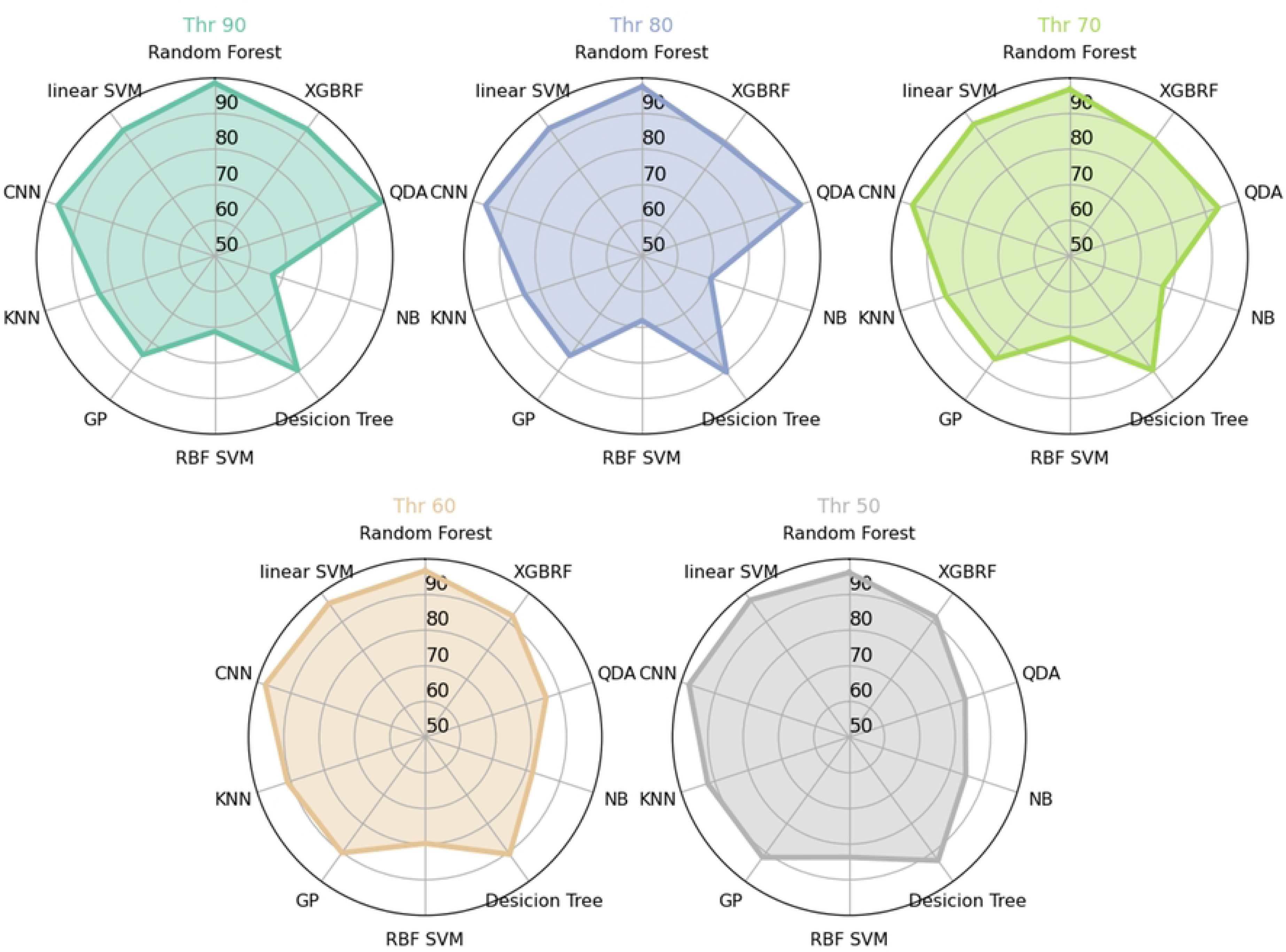
Performance of the classifiers. Visualization of the performance (accuracy) of the ten distinct classifiers on 2-second EEG epochs with 10,332 data points and 652 features in sensor space. Classifiers were applied using varying BATS thresholds from 90 to 50 with a step size of 10.

**Fig 3.**
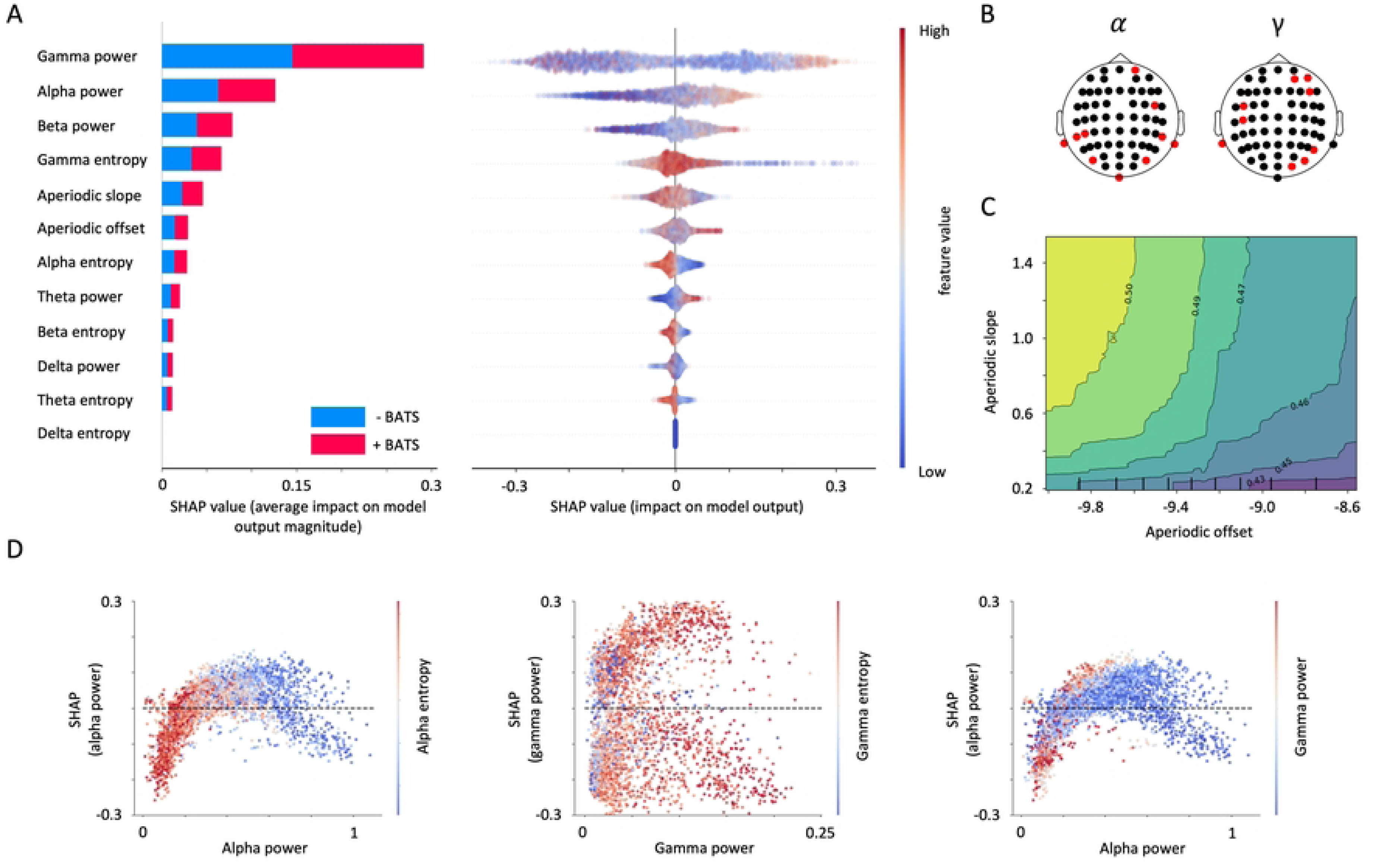
Importance order and direction of the sensor space features on predicting +BATS. **A**. On the left sub-panel, a list of sensor space features, arranged in decreasing order of importance in the classification procedure, is presented. The chart showcases the combined significance of features within each feature category for forecasting -BATS (absence of tinnitus acoustic suppression) and +BATS (presence of tinnitus acoustic suppression), indicated by blue and red colors correspondingly. The horizontal axis shows the averaged SHAP values associated with each feature category. Feature classes with higher average SHAP values have a greater impact on the prediction of targets. On the right sub-panel, the chart includes a collection of data points (i.e., single EEG epochs), which are placed horizontally across the x-axis, representing their respective SHAP values. Additionally, the color of each data point (epoch) reflects the feature values, with a gradient from red to blue representing high and low values. **B**. Channels of significance in the alpha (left panel) and gamma (right panel) frequency ranges are highlighted in red. **C**. Partial dependence plot displays the interaction between two feature sets (aperiodic offset and aperiodic slope) in predicting the class of individuals with +BATS. As the aperiodic offset decreases and the aperiodic slope increases, collectively indicating a reduction in non-oscillatory brain activity, there is a higher probability of predicting individuals with +BATS. **D**. The x-axis of the scatter plots represents the values of alpha power (left and right sub-panels) and gamma power features (middle sub-panel). Each data point corresponds to an individual observation (i.e., a single EEG epoch) in the dataset. The y-axis represents the SHAP values associated with each feature for the same set of data points, and color gradients represent a third variable, namely, alpha entropy (left panel), gamma entropy (middle sub-panel), and gamma power (left and right subpanel). The baseline (y = 0, dotted line) represents the model’s mean prediction of +BATS across all instances. Dots above the baseline indicate positive feature contribution, while dots below indicate negative feature contribution.

### Source space

Following the merging of features with high correlation, a total of 263 features, constituting 77% of the original 340, were excluded from our dataset. Subsequently, we trained the model and classified the data using these refined features, resulting in an accuracy of 97.8% for the RF model on the test data. Examining the overall importance of spectral power values within different frequency bands across brain labels indicated that delta, theta, and beta oscillations accounted for 10.3%, 13.3%, and 19.2% of the total feature importance, respectively. However, alpha and gamma oscillations contributed substantially more significantly, making up 27.2% and 29.8% of the total feature importance. This importance order aligns with results from the classification process performed in sensor space features. Fig 4 displays the key brain labels that contribute the most to the classification task, along with their predictive direction. The source model extends the findings of sensor space locations in the previous model by confining features to functionally-segregated brain regions. In general, alpha power was more pronounced in the right brain hemisphere whereas gamma power seems to exert a bias to the left hemisphere (Fig 4.A and B), at least regarding temporal and (primary) auditory fields. Furthermore, identified labels in primary auditory regions (transverse temporal, middle temporal) extend to non-auditory regions associated with sensory integration (superior parietal, supramarginal, precentral and paracentral), executive and attentional control (superior frontal, frontal pole), memory (parahippocampal, posterior cingulate), and limbic emotional (interface) integration (insula, rostral anterior cingulate, and temporal pole), for alpha and gamma, respectively. In the insula, high alpha power is not predictive of +BATS, whereas the opposite pattern can be observed for alpha power in the rostral anterior and the posterior cingulate cortex, and superior frontal gyrus. In the remaining labels of the alpha band, the impact on the model’s output is bidirectional or mixed. Gamma power is predictive of +BATS in (left) transverse temporal gyrus, rostral anterior cingulate cortex, and paracentral gyrus whereas it is not predictive in superior parietal and supramarginal gyri. For the remaining labels in gamma, their influence on the model’s output is observed to be bidirectional or variable.

**Fig 4.**
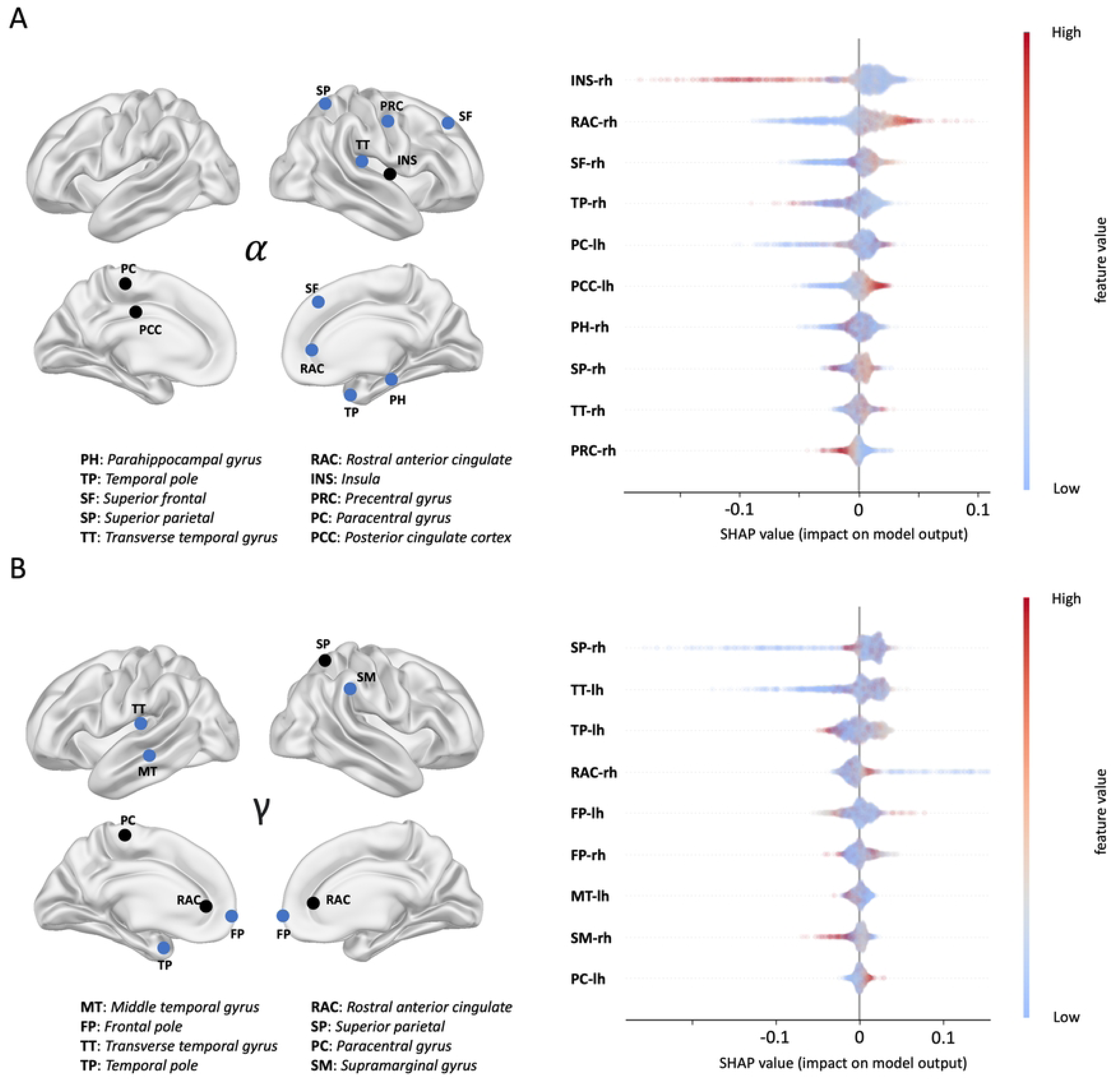
The most contributing brain labels in prediction of +BATS. In both the alpha (**A**) and gamma (**B**) frequency range, brain labels that significantly contributed to predicting individuals with +BATS are denoted by circles. Brain labels shared with the validation dataset are highlighted in blue, while those not matching are colored in black. The importance and directionality of these brain labels in predicting +BATS are displayed in the right subpanels of **A** and **B**, respectively.

### Connectivity

We calculated the coherence connectivity measure within the resting state network’s brain labels and between the important brain labels identified in the source space analysis. The coherence measures were computed for each epoch, and 20 features out of 102 features were excluded due to high correlations (19.6%). Consequently, the training phase involved 82 features, encompassing connections between resting state networks and auditory network brain labels within two frequency ranges, specifically alpha and gamma. After training the model, we achieved an accuracy of 86.3%. The most important connections for both frequency ranges are presented in Fig 5. Overall, the predictive feature set of this model is driven by important gamma connections between several networks and nodes while important connectivity in the alpha frequency band was limited to 3 between-network connections and 2 intra-auditory network connections (hyperconnectivity between bilateral primary auditory fields in superior and transverse temporal gyrus as well as superior parietal gyrus. Interestingly, all alpha between-network connections (i.e., VAN and DMN or DGN, AUN and DAN) were not predictive of +BATS indicating a global and trait-like decoupling of these networks +BATS individuals. In contrast, intra-auditory network connections in the alpha band (i.e., between superior temporal, parietal, and transverse temporal gyri) are predictive of +BATS. Gamma connectivity predictive of +BATS resulted between AUN and SMN, AUN and LBN, AUN and DGN, SMN and FPN, and VAN and DAN, while the remaining connectivity features had mixed or negative impact on the model output (i.e., VAN and DGN, AUN and FPN, VAN and LBN, and VAN and SMN). Finally, a single intra-auditory network connection in the gamma frequency band predictive of +BATS was found between left superiorparietal gyrus and right superiortemporal gyrus.

**Fig 5.**
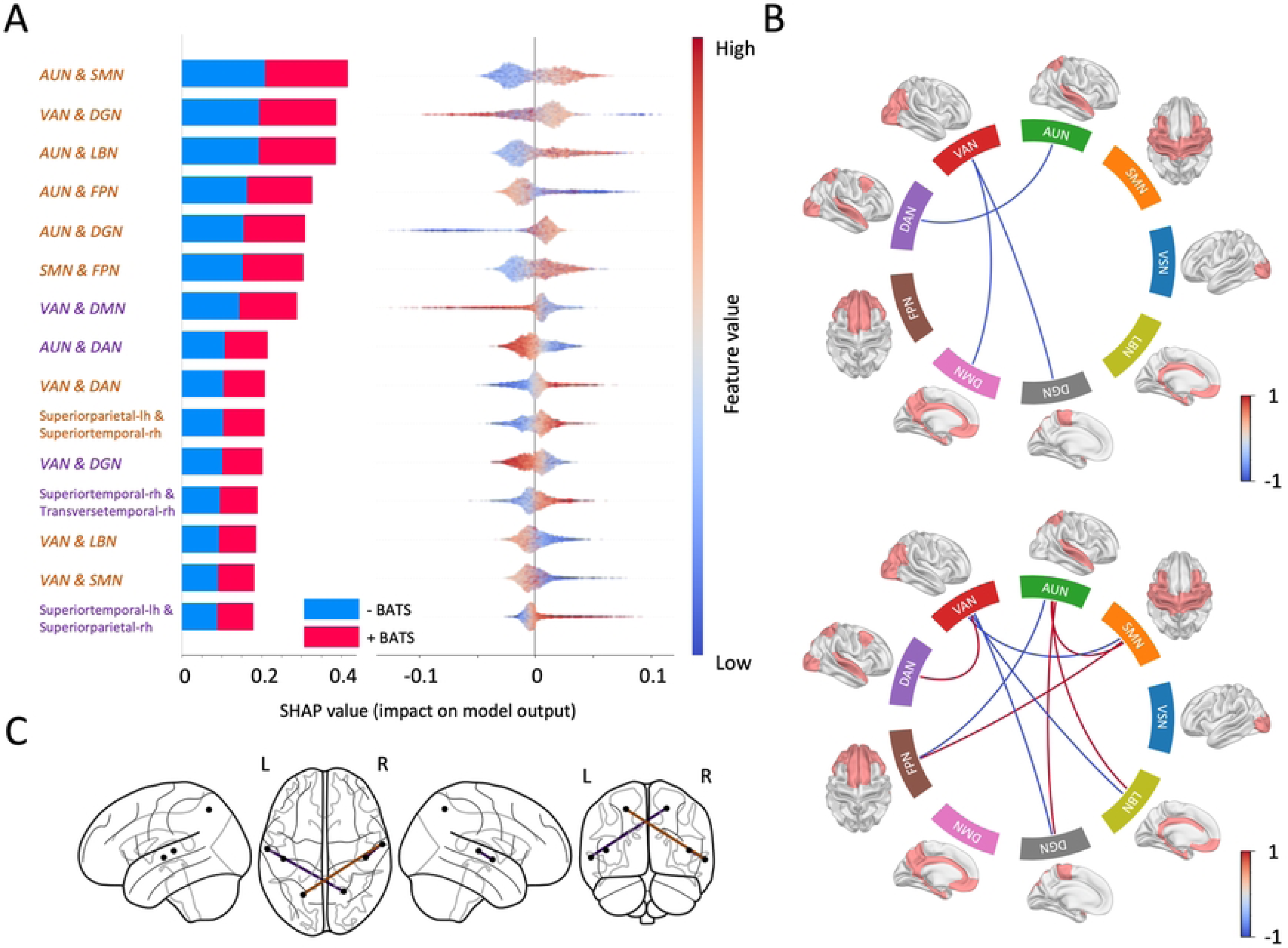
Importance order and direction of the connectivity features on predicting +BATS. **A**. The list of connections, sorted by importance in the classification process along with their directionality in predicting +BATS and -BATS, is presented. The y-axis tick labels, colored in purple and light brown, represent connections in the alpha and gamma frequency ranges, respectively. **B**. The contrast between average connectivity across participants with +BATS and -BATS in alpha (upper panel) and gamma (bottom panel) within resting-state networks is depicted using circular graphs. The color gradient ranges from red to blue, indicating stronger connections in individuals with BATS ability to low values, indicating individuals with -BATS. **C**. The most important connections within the AUN network in the classification process are visualized on a glass brain. Purple and light brown indicate connections in the alpha and gamma frequency ranges, respectively.

### Validation

We conducted an ancillary analysis using a validation dataset to assess the generalizability of our findings and whether the identified important brain labels were consistent across different recording systems (see S6 Fig and S7 Fig) and varied levels of +BATS obtained from different response scales (see S5 Fig). Spectral power values were computed in brain parcels, following the same methodology outlined in section Feature extraction. Feature merging was performed to address high correlations, resulting in 107 (68.5%) features being retained from the initial 340 features and data classification was carried out in the sensor space, using the remaining features with a selected loudness threshold value of −1. An RF model was employed, and an accuracy of 99.1% was achieved on the test data, using the top 100 most important features. When comparing the important brain labels obtained from the validation dataset analysis with the top 10 most important brain labels from the main dataset, we observed that in the alpha frequency range, 7 out of the 10 brain labels were important in both datasets. The 3 non-overlapping brain labels were the paracentral gyrus, insula, and posterior cingulate. Notably, the reason for their absence in the validation set was the presence of high correlations with other brain parcels, which led to their removal in the initial feature merging step. Specifically, the paracentral gyrus exhibited high correlation with labels: postcentral, posterior cingulate, superior parietal, and supramarginal. The insula displayed high correlation with brain labels: lateral orbitofrontal, medial orbitofrontal, pars opercularis, pars orbitalis, pars triangularis, rostral middle frontal, and superior temporal, and the posterior cingulate was removed due to its high correlation with the supramarginal label. In the gamma frequency range, 7 out of the 10 most important brain labels matched between the two datasets. Similarly, 3 out of the 10 brain labels did not appear in the validation dataset. These were the rostral anterior cingulate (due to high correlation with superiortemporal and temporal pole), paracentral (due to high correlation with postcentral, posterior cingulate, superior parietal, and supramarginal), and superior parietal (due to high correlation with supramarginal).

Finally, we conducted training and testing of a RF model on the validation data using connectivity features computed as described in Section Feature extraction. This analysis resulted in an accuracy of 82.2%. When we compared the most important connections derived from the main dataset with those from the validation dataset, we found that most of the important connections were present in the validation dataset’s results, while both the order and the extent of the feature list vary considerably. In general, considering that there is a 70% match between the important brain labels in both the alpha and gamma frequency ranges and taking into account that the remaining labels were dropped due to the feature merging process, we can conclude that the validation dataset successfully validated the findings from the main dataset. This level of consistency supports the robustness and reliability of our results across different datasets and recording systems.

## Discussion

In the present study, we aimed to ascertain if specific sensor, source, or connectivity features of resting state EEG from individuals with tinnitus predict tinnitus suppression by auditory stimulation. We showed that high classification accuracy can be found for several BATS threshold levels (split validation) and in an independent dataset. Important neural features were identified and subjected to model impact and directionality (of effects) analysis, which resulted in specific patterns of neural signatures aligning and extending current models of tinnitus. In the absence of any directly comparable previous work (i.e., prediction of acoustic tinnitus suppression from naive (EEG) resting state data and not by experimental state-like BATS data), referential discussion within the tinnitus literature in the following section is inherently limited. We first discuss the classification workflow and results, followed by an integrative discussion of resulted neural features with respect to tinnitus and general brain models following the sequence of analysis steps (see Fig 1). Limitations, future directions, and the conclusion will finally complement the discussion.

### Classification

Our analysis demonstrated the feasibility of robustly classifying individuals with regards to acoustically-induced tinnitus suppression based on naive EEG resting state recordings. We demonstrated that the classification task remains robust and consistently yields high accuracy on unseen data for various BATS threshold values (see Fig 2). This was further confirmed by our ancillary randomly-shuffled label model analysis, which resulted in almost chance-level accuracy (51.7%). In contrast to classical approaches that focus primarily on designing classifiers, even complex ones, to achieve high accuracy without a deep exploration of the underlying dynamics, the emphasis of this work is distinct: The focus lies not in classification per se, but in meaningful, explainable outcomes that foster the understanding of the underlying patterns. Showing that various simple models consistently yield high classification accuracy implies that the problem inherently possesses a global minimum in the parameter space for all classifiers. This implies that various simple models converge to the same optimal solution, indicating robustness and reliability across different approaches to classification. Beyond that, we assessed the importance and directionality of the feature classes for different loudness threshold levels, as illustrated in supplementary Fig S5 Fig. This ancillary analysis further highlighted consistency in both importance and direction across the class of features. Any (usually small) variations observed may be mostly attributed to the random selection of training and test sets, shuffling or data imbalance. This suggests that the choice of the threshold does not strongly influence the data’s underlying pattern. This robustness indicates that the features have been well-designed and offer clear separability between the two classes, resulting in consistent model performance across different thresholds. The ultimate choice of a 10% BATS threshold (tinnitus loudness threshold at 90% after stimulation) for the main analysis is thus solely motivated to create a balanced dataset. This threshold results in a distribution where 47.9% of individuals have +BATS, and 52.1% have -BATS, which ultimately reduces model bias and leads to fairer predictions.

Since preserving the inherent meaning of our features is important, we did not employ advanced feature selection or reduction techniques like PCA [59] or NCA [60], which involve linear combinations of features. We instead utilized Pearson correlation approach to merge features that exhibit high correlations to address the issue of multicollinearity in our dataset while keeping features interpretable. This process reduced dataset dimensionality, eliminated multicollinearity, and mitigated overfitting, retaining essential data for classification. However, features that exhibit a non-linear connection may be retained despite potentially showing a weak Pearson correlation, as the Pearson correlation specifically assesses linear associations. Merging such features based on correlation could be detrimental in certain configurations. Moreover, choosing the wrong threshold for merging the features can lead to either under-merging (retaining too many features) or over-merging (losing important information).

We ultimately selected the RF model to classify individuals with respect to their ability to suppress their tinnitus for several reasons: First, its majority voting mechanism naturally mitigates the risk of overfitting and helps lower variance error, promoting more robust and reliable predictions [48] (see supplementary table S4 Table). Second, RF is known for being less sensitive to hyperparameter choices compared to other models. Third, it offers a measure of feature importance through the Gini impurity metric [61]. Additionally, it consistently demonstrated superior performance across various loudness thresholds, further validating its suitability for the task. Lastly, RF is an ensemble which is helpful if unbalanced data is present, in contrast to other classifier methods. Therefore, its inclusion and application on different BATS splits can be considered as optimal.

Features with low importance may have limited impact on the model’s predictive capabilities and can potentially be removed to simplify the interpretation of the model. On the other hand, if feasible and of interest, one could consider their effect on the model’s output as well. As an example, when examining non-oscillatory features of power spectrum density, including PSD offset and slope, we observed that as moving towards lower offset values and higher slope values (indicating reduced aperiodic activity), the model tends to predict the class of individuals who consistently showed tinnitus suppression (+BATS). This relationship is illustrated in Fig 3.C. Taken together, the proposed and applied classification method seems to be very feasible to investigate trait-like neural features with regards to their predictive value on the ability to (acoustically) suppress tinnitus.

### Neurophysiological relevance

#### Sensor space

In the first segment of our discussion, we focus on the model and features regarding the EEG sensor space. First, it’s worth noting that higher gamma power values were more predictive of +BATS, with a distinct bias towards positive predictions but some negative extremes as well. In the absence of directly comparable experimental data, the discussion here is limited to links to general resting state data of tinnitus (trait-like) and to links to neural signatures during BATS (state-like). The latter comparison is especially precarious given the absence of state-like data and results in our study. Yet, looking at the pattern of positively-biased but bidirectional pattern of gamma features, our analysis aligns with the findings of Sedley et al. [31], where they argued that there was a positive correlation between tinnitus intensity and gamma band oscillations in the auditory cortex among a majority of patients (8 out of 14), suggesting an increased thalamocortical input and cortical gamma response associated with higher tinnitus loudness. Conversely, all four patients exhibiting residual excitation (i.e., tinnitus loudness exceeding the baseline loudness before sound stimulation) demonstrated an inverse correlation between perceived tinnitus intensity and auditory cortex gamma oscillations. In a further study, it was shown that gamma oscillations are consistently more present during BATS [32], which was recently confirmed by another study [35], where an increase in alpha and gamma frequency bands during BATS was shown. Contrary to these positive gamma findings, two other studies could not find gamma and/or high frequency oscillation effects in BATS [34, 62]. In our previous study, we observed decreased low gamma or high beta power post-stimulation (at 31 Hz), which was not linked to BATS [36]. The BATS experimental data regarding gamma thus seems to be inconclusive and, as introduced, considering general resting state data from tinnitus and basic literature about gamma or neural oscillations might be more productive. In comparison to healthy controls, increased gamma power in rest in individuals with chronic subjective tinnitus has been found in several studies [12–14, 63]. In addition, some resting-state studies have shown a positive correlation between tinnitus loudness and gamma oscillations in auditory fields [64, 65], which was also critically discussed in a position paper [66]. The pattern of increased high-frequency or gamma oscillations in tinnitus resting-state seems to be stable, with little contradicting evidence, and may be interpreted with the theorized higher neural synchrony in cortical auditory fields due to tinnitus [9, 67]. Mapping these considerations back to our novel data, we certainly can assume that the gamma findings, including some bidirectional effects possibly related to individual differences, are reflected in our findings. Yet, it is not understood in detail how gamma oscillations may contribute to an active suppression of tinnitus. Gamma oscillations, typically in the range of 30-100 Hz, are closely associated with sensory processing, attention, and the integration of cortical information [68, 69]. In the context of the auditory system, higher gamma power could reflect enhanced cortical excitability and increased neural synchrony within auditory pathways, which might be instrumental in the modulation or suppression of tinnitus. We thus theorize that increased trait-like gamma power and/or dynamics, as found in our study, might be aiding the cortical (auditory) system to suppress tinnitus. In addition, current considerations regarding the predictive brain might extend this reasoning by introducing that higher gamma activity in auditory cortical fields could be interpreted as the brain’s attempt to enhance the precision of auditory predictions or to amplify the prediction error related to the phantom sound of tinnitus [32, 70, 71]. This increased activity could serve to better predict and, therefore, more effectively cancel out the internal representation of tinnitus, leading to its suppression.

Second, looking at the second most important identified feature, alpha power, the discussion of our finding of increased alpha power predictive of +BATS is more straightforward both given the clear direction of results and the results’ fit to current theoretical models. Reduced (trait-like) alpha power in tinnitus has been consistently shown in resting state studies [12, 72] and is theorized to be reflective of a disrupted (auditory) inhibitory system in tinnitus. Findings of reduced GABA, a major inhibitory neurotransmitter, concentration levels in cortical auditory fields may further corroborate the hypothesis of a defective inhibitory system in tinnitus [73, 74]. A single study also showed that observed lower resting alpha power in tinnitus is correlated to higher gamma power linking the two major inhibitory and excitatory neural oscillations [63]. We could demonstrate a similar correlation in our analysis of +BATS prediction (see Fig 3.D). Furthermore, experimental data in BATS showing increases of alpha power during BATS confirms that BATS may temporarily restore normal cortical inhibition and thus suppress the perceived tinnitus sound [75]. Taken together, our results add to the importance of alpha regarding cortical inhibition and (acoustic) tinnitus suppression by establishing its importance as a trait-like feature in individuals with tinnitus, which has not been shown before.

Finally, aperiodic parameters complemented the results of the sensor feature set, demonstrating that +BATS is related to lower aperiodic offsets and steeper slopes, which suggest more periodic (i.e., oscillatory) activity over the entire power spectrum in individuals with +BATS (Fig 3.C). This observation fits considerations of thalamo-cortical dysrhythmia in tinnitus [76–78], expressed by a flatter overall M/EEG (resting state) spectrum and increases in high-frequency power (i.e., gamma).

#### Source space

Source localization of the identified most important features of the sensor model, gamma and alpha power, was motivated to constrain these findings to specific functionally-segregated brain regions. The discussion so far assumed that the effects to be mainly originating from cortical (bilateral) auditory fields, which is confirmed by the resulting sensor locations spanning lateral and posterior sensors (see Fig 3.D) and the bulk or previous literature. Regarding frequency-specific contributions in identified brain labels, global functional eminence of the alpha and gamma band in the context of tinnitus and sensory processes have to be elucidated: The prominence of alpha power in the right hemisphere and gamma in the left hemisphere suggests a division of labor between the hemispheres in general brain functioning [79]. Alpha oscillations generally reflect inhibitory processes and reduced cortical arousal [80], whereas gamma oscillations typically code sensory processing, attention, and the integration of cortical information [69]. Mapping these global and normal brain mechanisms to tinnitus and especially to trait-like features predicting tinnitus suppression is challenging in the absence of any relevant previous resting state data in tinnitus considered with lateralization of neural oscillations. Yet, our findings suggest that there is a correlation between the level of predictive gamma power in the left transverse temporal gyrus and predictive alpha power in the right transverse temporal gyrus (as shown in Fig 4.A and B), and the typical functioning preferences of the two hemispheres, particularly in the bilateral auditory cortex. This implies that normal brain functioning may be facilitating (acoustic) tinnitus suppression. Predictive trait-like gamma power in the left primary auditory cortex, namely transverse temporal or Heschl gyrus, may signify an adaptive neural process aimed at minimizing the sensory prediction errors that underlie the perception of tinnitus. In turn, such a minimization of prediction errors may contribute to successful tinnitus suppression.

In the insula, high alpha power is not predictive of +BATS, whereas the opposite pattern can be observed for alpha power in the rostral anterior cingulate cortex, posterior cingulate cortex, and superior frontal gyrus. In the rostral anterior cingulate cortex, predictive trait-like alpha power could be linked to a functioning or maintained active tinnitus noise canceling system mediated through thalamocortical relays [81]. A correlative EEG resting state study has identified the anterior cingulate complex, including the rostral anterior cingulate cortex, to be involved in tinnitus perception [82]. Yet, the authors did not find any effects in the alpha frequency band. Moreover, also critical for our data, M/EEG source localization of deep, medial, and ventral structures like the subcallosal area, identified as the key node in the tinnitus noise-cancelling system [81, 83], is challenging and possibly unreliable. Given structural vicinity and functional overlap of the anterior cingulate complex’ subregions (i.e., rostral anterior cingulate cortex, pregenual anterior cingulate cortex, and subgenual anterior cingulate cortex), it was proposed by [84] to extend the functional locus of the key node of the noise-canceling system to the entirety of the ventromedial prefrontal cortex. Predictive trait-like alpha power in the posterior cingulate cortex could be both reflective of an intact DMN including its inhibitory properties and/or normal modes of memory processing, which could imply that tinnitus is not filled-in from the hippocampus as proposed in recent models [85, 86]. In similar veins, predictive alpha in (superior) frontal regions could be indicative of functioning control (networks) within the tinnitus brain, allowing for better attentional (or auditory gating) control and possibly suppression of the phantom sound perception [87]. Looking at gamma, its lack in superior parietal regions may be correlated to an absence of integration with other sensory systems (cross-modal compensation) and/or intact attentional or inhibitory control as in alpha and the posterior cingulate cortex [88, 89]. Finally, the same could be true for a similar pattern in the rostral anterior cingulate cortex and, analogously to higher predictive alpha power in the same region, linked to an (intact) tinnitus noise canceling system.

Notably, identified regions also play a role in large and small-scale networks, such as, for example, the posterior cingulate cortex, a critical node in the default mode network. The involvement of the insula and the rostral anterior cingulate cortex highlights their significance in the salience network or ventral attention network, crucial for detecting and filtering salient external stimuli and internal events, thereby facilitating the transition between activated networks such as the default mode network and the central executive or frontoparietal network. The superior frontal gyrus and the frontal pole, implicated in executive function and attentional control, are key components of the frontoparietal network, underscoring their role in directing attention and managing cognitive resources, which could be particularly relevant in modulating attention towards or away from tinnitus sounds [90]. Additionally, the involvement of sensory integration and executive regions suggests a complex interplay between auditory processing and higher-order cognitive functions, emphasizing the multisensory and multidimensional nature of tinnitus perception within these overarching neural networks. Network aspects are further analyzed and discussed in the following section discussing our network model.

#### Connectivity

Looking at auditory connectivity, the specific intra-auditory alpha and gamma connections in +BATS individuals most probably reflect functioning inhibitory circuitry enabling the suppression of tinnitus (see Fig 5.C. The here observed intact connectivity contradicts findings of disrupted resting state alpha networks in individuals with tinnitus [91], implying that an intact intra-auditory network may support BATS. In the same study, authors found a resting state hyperconnection in the gamma frequency range which could be explained by differences in the analysis (i.e., resting state case-control design in the former study vs. within-group prediction in our study). In general, the resting state auditory network literature in tinnitus is not unequivocal, with conflicting results explained by the heterogeneity of the investigated tinnitus samples and/or applied methods [16].

On the large-scale network level, in the gamma frequency band, the most important connection of the connectivity RF model is found between AUN and SMN possibly related to increased sensory integration and/or attempts to minimize sensory prediction errors. Notably, both networks include the transverse temporal gyrus, which highlights their intrinsic connection and, thus, functional coupling. Further, a hyperconnection between AUN and the LBN could be representative of a trait-like intact noise canceling system [81], which may be further corroborated by a similar hyperconnection between AUN and the DGN including bilateral caudate, putamen, pallidum, and thalamus. However, given the large-scale character of investigated networks, more precise identification of medioventral key nodes of the proposed noise-canceling system (i.e., subgenual cingulate cortex and/or anterior cingulate cortex complex as well as the thalamus), can not be provided with our current data and analyses. Contrary to these hyperconnections, a hypoconnection between AUN and the (control) network FPN could be characteristic of a trait-like pattern of less attentional control and/or memory-related connectivity in +BATS individuals. This could imply that individuals with +BATS may not have developed aberrant network activity. The observed 3 hypoconnections between large-scale networks in the alpha frequency band, namely between VAN and DMN, AUN and DAN, and VAN and DGN, respectively, may echo the hypoconnection between AUN and FPN in the gamma frequency band. This would further corroborate the emerging pattern of hyperconnected auditory and/or potential noise-canceling networks in the absence of interactions between other large-scale networks and/or with the hyperconnected networks predicting +BATS.

Taken together, connectivity results of our 3rd model unfathomed a global pattern of intact intra-auditory connections in both frequency bands possibly implying functioning inhibitory and/or integrating auditory circuitry. Beyond that, large-scale networks are mostly hypoconnected, except auditory sensory as well as auditory limbic interactions, indicating normal functioning and/or an unimpaired noise-canceling system.

#### Behavioral differences

In our analysis, we observed higher MML and a trend towards higher tinnitus loudness levels in the -BATS group, suggesting a potential relationship between tinnitus perceptual intensity and the ability to achieve BATS. This observation is intriguing, especially given the absence of significant differences in tinnitus duration between groups, a marker often associated with tinnitus chronification. By crossing a non-linear threshold where the system’s adaptive responses may become maladaptive, certain inhibitory and/or neuroplastic mechanisms necessary for tinnitus suppression may become less effective [92, 93]. This observation points to a need for further investigation into the neural and perceptual dynamics that limit tinnitus modulation.

### Limitations and future directions

The current study has some limitations that inform future studies in BATS, tinnitus research, and/or the methods applied.

Classifier feature importance values and related ordered lists are delicate to interpret. Resulting values are not straightforward to interpret per se, especially if compared to modes of interpreting inferential frequentist or Bayesian statistics results. In a RF classifier, the Gini importance values, which quantify each feature’s contribution to node purity and the overall quality of splits across the decision trees, do not necessarily sum to 1 but are scaled relative to each other. These values, reflective of a feature’s frequency in splitting and its impact on reducing node impurity, vary with the data, the forest’s size, and the algorithm’s implementation, allowing for a meaningful comparison of feature importance within the model’s context. SHAP values are used to explain the output of machine learning models by attributing the prediction to different features. They represent the contribution of each feature to the difference between the actual prediction and the average prediction. Each SHAP value corresponds to a feature and indicates how much that feature contributes to the prediction for a particular instance. Despite the complexities inherent in interpreting Gini importance and SHAP values within a RF classifier, our study’s results remain reliable and interpretable, acknowledging the discussed limitations.

Even though our results are of high accuracy, stable over validation approaches, and meaningful in resulting features, larger sample sizes are needed to further consolidate and differentiate analyses and results. Yet, given our total sample size of 102 cases, the presented dataset is currently the largest in the context of EEG, BATS, and tinnitus.

Further, future studies could incorporate additional neurophysiological measures of higher spatial resolution, such as MRI, to complement the current feature sets and to ensure more precise source localization based on individual structural MRI in combination with scanned individual EEG electrode positions. Source localization may limit the precision of some of the presented data, which is discussed transparently throughout the paper.

To maximize the feature set included in machine learning modeling and, in consequence, derive insights in the spirit of explainable AI [37], further (neuro)physiological measures could be considered [94]. A maximized comprehensive feature set could lead to objective diagnosis and subtypization of tinnitus and/or the ability of tinnitus suppression [95, 96].

Finally, by mapping the unique neural signatures associated with tinnitus in different individuals derived from our approach here, future studies could design targeted interventions that address the specific neural underpinnings of tinnitus in each individual. Such an approach would not only improve the precision of tinnitus treatments but also contribute to the broader field of personalized neurotherapy, optimizing interventions based on each individual’s neural fingerprint.

## Conclusion

The present work represents the first attempt to predict acoustic tinnitus suppression via spontaneous brain activity data. It aims to understand the potential suppression factors on the neural level through automatic classification and identification of distinctive features. We achieved high classification accuracy (98% for the sensor and source model and 86% for the connectivity model) and identified several partly novel, trait-like neural features critical for understanding tinnitus suppression. By revealing specific patterns of gamma and alpha oscillations in sensors and specific brain regions, our results extend current models of tinnitus, highlighting the role of auditory cortical activity and its hemispheric distribution in managing phantom sound perception and suppression. Furthermore, we could demonstrate that intra-auditory and cross-network connectivity between large-scale (cortical) auditory and limbic networks were also predictive of an individual’s ability to suppress tinnitus. Finally, by analyzing aperiodic features of the EEG power spectrum, it was shown that normal averaged spectral shapes are predictive of tinnitus suppression. Our approach not only advances the comprehension of the neural basis of tinnitus suppression and tinnitus in general but also paves the way for objective diagnosis and personalized treatment strategies.

## Supporting information

**S1 Table. Sample description.** Information about the samples, including sex, tinnitus side, age, tinnitus duration, hearing loss, MMl, THI score, GUF score, tinnitus loudness, and BATS% is presented for both the main and validation datasets.

**S2 Table. Descriptive statistics of sample split in the main dataset.**

**S3 Table. Overview of brain regions extracted from Desikan-Killiany atlas and assigned to canonical 9 sub-networks.**

**S4 Table. Random Forest classifier performance.** Metrics evaluating performance, such as accuracy, precision, recall, and F1-score, are provided for each classification task, encompassing both the primary dataset and the validation dataset.

**S5 Fig. Importance order and directionality of sensor space features are shown for three different loudness threshold value, namely 90 (A), 70 (B) and 50 (C).** The dots at each plot indicate SHAP values measuring how much each feature category contributes to predicting class of individuals with +BATS. Furthermore, the color of each data point (epoch) represents the feature values, following a gradient from red to blue, where red indicates high values, and blue signifies low values.

**S6 Fig. Analysis of validation dataset. A**. Visual representation of the correlation matrix of the validation dataset, showing the relationships between various pairs of features including computed features in sensor space (left panel), source space (middle panel) and connectivity features (right panel). To reduce multicollinearity among features, values greater than 0.9 are combined. **B**. Accuracy values of ten distinct classifiers applied on the EEG epochs of validation dataset. **C**. Sensor space features are organized by decreasing importance for predicting two classes: +BATS and -BATS. In the left panel, the horizontal axis displays averaged SHAP values for each feature category, with higher values indicating greater influence on target prediction. Right panel consists of data points (epochs), representing feature categories, placed along the x-axis based on their SHAP values. Data point colors range from red to blue, reflecting feature values from high to low. **D**. Most contributing channels in classifying individuals with +BATS and -BATS are colored in red in both alpha (right panel) and gamma (left panel) frequency ranges.

**S7 Fig. The list of most contributing connections in the validation dataset, sorted by their importance in the classification process.** Each connection’s directionality is also shown. The y-axis tick labels, colored in purple and light brown, correspond to connections in the alpha and gamma frequency ranges, respectively.

**S8 Text. Supplemental information.** This section outlines the EEG recording procedure and data preprocessing pipeline. Furthermore, techniques for extracting spectral band powers, spectral entropy, aperiodic spectral power, source space power spectral density, and connectivity measures are described. Additionally, methods for evaluating the importance and directionality of features are introduced.

## Acknowledgments

We are thankful to Susanne Staudinger, Anita Hafner, Bernhard Unsin, Kira Voigt, Jonathan Kisskalt, Anthony Ngu, Mariana Martins-Lopez, Deniza Avdi, Vithushika Raveenthiran, and Allegra Preisig for their invaluable help in study management and data acquisition. We thank all the participants for their efforts and patience in the partly tedious experimental procedures. Payam S. Shabestari and Patrick Neff have been funded by SNF (Project Grant number 325130 208164) for this work.

## Author Contributions

**Conceptualization:** Payam S. Shabestari, Stefan Schoisswohl, Martin Schecklmann, Patrick Neff

**Data Curation:** Payam S. Shabestari, Stefan Schoisswohl, Zino Wellauer, Patrick Neff

**Formal analysis:** Payam S. Shabestari, Stefan Schoisswohl, Patrick Neff

**Funding acquisition:** Tobias Kleinjung, Berthold Langguth, Patrick Neff

**Investigation:** Stefan Schoisswohl, Zino Wellauer, Patrick Neff

**Methodology:** Payam S. Shabestari, Patrick Neff

**Project administration:** Patrick Neff

**Resources:** Tobias Kleinjung, Martin Schecklmann, Berthold Langguth

**Supervision:** Tobias Kleinjung, Martin Schecklmann, Berthold Langguth, Patrick Neff

**Writing - Original Draft:** Payam S. Shabestari, Patrick Neff

**Writing - Review & Editing:** Stefan Schoisswohl, Zino Wellauer, Adrian Naas, Tobias Kleinjung, Martin Schecklmann, Berthold Langguth

## References

1. De Ridder D, Schlee W, Vanneste S, Londero A, Weisz N, Kleinjung T, et al. Tinnitus and tinnitus disorder: Theoretical and operational definitions (an international multidisciplinary proposal). 2021;260:1–25. doi:10.1016/bs.pbr.2020.12.002.

2. Baguley D, McFerran D, Hall D. Tinnitus. The Lancet. 2013;382(9904):1600–1607. doi:10.1016/S0140-6736(13)60142-7.

3. Jarach CM, Lugo A, Scala M, van den Brandt PA, Cederroth CR, Odone A, et al. Global Prevalence and Incidence of Tinnitus: A Systematic Review and Meta-analysis. 2022;79(9):888–900. doi:10.1001/jamaneurol.2022.2189.

4. Pinto PCL, Marcelos CM, Mezzasalma MA, Osterne FJV, de Melo Tavares de Lima MA, Nardi AE. Tinnitus and its association with psychiatric disorders: systematic review. 2014;128(8):660–664. doi:10.1017/S0022215114001030.

5. Langguth B, Kleinjung T, Schlee W, Vanneste S, De Ridder D. Tinnitus Guidelines and Their Evidence Base. 2023;12(9):3087. doi:10.3390/jcm12093087.

6. Simoes JP, Daoud E, Shabbir M, Amanat S, Assouly K, Biswas R, et al. Multidisciplinary Tinnitus Research: Challenges and Future Directions from the Perspective of Early Stage Researchers. 2021;13. doi:10.3389/fnagi.2021.647285.

7. McFerran DJ, Stockdale D, Holme R, Large CH, Baguley DM. Why Is There No Cure for Tinnitus? 2019;13. doi:10.3389/fnins.2019.00802.

8. Langguth B, Kreuzer PM, Kleinjung T, De Ridder D. Tinnitus: causes and clinical management. Lancet Neurol. 2013;12(9):920–930. doi:10.1016/S1474-4422(13)70160-1.

9. Eggermont JJ, Tass PA. Maladaptive Neural Synchrony in Tinnitus: Origin and Restoration. Front Neurol. 2015;6. doi:10.3389/fneur.2015.00029.

10. Elgoyhen AB, Langguth B, De Ridder D, Vanneste S. Tinnitus: perspectives from human neuroimaging. Nature Reviews Neuroscience. 2015;16(10):632–642. doi:10.1038/nrn4003.

11. Eggermont JJ, Roberts LE. The Neuroscience of Tinnitus: Understanding Abnormal and Normal Auditory Perception. Front Syst Neurosci. 2012;6. doi:10.3389/fnsys.2012.00053.

12. Weisz N, Moratti S, Meinzer M, Dohrmann K, Elbert T. Tinnitus Perception and Distress Is Related to Abnormal Spontaneous Brain Activity as Measured by Magnetoencephalography. PLoS Med. 2005;2(6):e153. doi:10.1371/journal.pmed.0020153.

13. Weisz N, Müller S, Schlee W, Dohrmann K, Hartmann T, Elbert T. The neural code of auditory phantom perception. J Neurosci. 2007;27(6):1479–1484. doi:10.1523/JNEUROSCI.3711-06.2007.

14. Ashton H, Reid K, Marsh R, Johnson I, Alter K, Griffiths T. High frequency localised “hot spots” in temporal lobes of patients with intractable tinnitus: A quantitative electroencephalographic (QEEG) study. Neurosci Lett. 2007;426(1):23–28. doi:10.1016/j.neulet.2007.08.034.

15. Vanneste S, Heyning PVd, Ridder DD. Contralateral parahippocampal gamma-band activity determines noise-like tinnitus laterality: a region of interest analysis. Neuroscience. 2011;199:481–490. doi:10.1016/j.neuroscience.2011.07.067.

16. Kok TE, Domingo D, Hassan J, Vuong A, Hordacre B, Clark C, et al. Resting-state Networks in Tinnitus. Clinical Neuroradiology. 2022; p. 1–20. doi:10.1007/s00062-022-01170-1.

17. Piarulli A, Vanneste S, Nemirovsky IE, Kandeepan S, Maudoux A, Gemignani A, et al. Tinnitus and distress: an electroencephalography classification study. Brain Communications. 2023;5(1):fcad018. doi:10.1093/braincomms/fcad018.

18. Jianbiao M, Xinzui W, Zhaobo L, Juan L, Zhongwei Z, Hui F. EEG signal classification of tinnitus based on SVM and sample entropy. Computer Methods in Biomechanics and Biomedical Engineering. 2023;26(5):580–594. doi:10.1080/10255842.2022.2075698.

19. Hong ES, Kim HS, Hong SK, Pantazis D, Min BK. Deep learning-based electroencephalic diagnosis of tinnitus symptom. 2023;17:1126938. doi:10.3389/fnhum.2023.1126938.

20. Allgaier J, Neff P, Schlee W, Schoisswohl S, Pryss R. Deep Learning End-to-End Approach for the Prediction of Tinnitus based on EEG Data*. 2021 43rd Annual International Conference of the IEEE Engineering in Medicine & Biology Society (EMBC). 2021;00:816–819. doi:10.1109/embc46164.2021.9629964.

21. Neff P, Michels J, Meyer M, Schecklmann M, Langguth B, Schlee W. 10 Hz Amplitude Modulated Sounds Induce Short-Term Tinnitus Suppression. Front Aging Neurosci. 2017;9. doi:10.3389/fnagi.2017.00130.

22. Neff P, Zielonka L, Meyer M, Langguth B, Schecklmann M, Schlee W. Comparison of Amplitude Modulated Sounds and Pure Tones at the Tinnitus Frequency: Residual Tinnitus Suppression and Stimulus Evaluation. Trends Hear. 2019;23:2331216519833841. doi:10.1177/2331216519833841.

23. Schoisswohl S, Arnds J, Schecklmann M, Langguth B, Schlee W, Neff P. Amplitude Modulated Noise for Tinnitus Suppression in Tonal and Noise-Like Tinnitus. Audiol Neurotol. 2019;24(6):309–321. doi:10.1159/000504593.

24. Tyler R, Stocking C, Secor C, Slattery WH. Amplitude Modulated S-Tones Can Be Superior to Noise for Tinnitus Reduction. Am J Audiol. 2014;23(3):303. doi:10.1044/2014*_A_JA* − 14 − 0009.

25. Reavis KM, Rothholtz VS, Tang Q, Carroll JA, Djalilian H, Zeng FG. Temporary Suppression of Tinnitus by Modulated Sounds. J Assoc Res Otolaryngol. 2012;13(4):561–571. doi:10.1007/s10162-012-0331-6.

26. Fournier P, Cuvillier AF, Gallego S, Paolino F, Paolino M, Quemar A, et al. A New Method for Assessing Masking and Residual Inhibition of Tinnitus. Trends Hear. 2018;22. doi:10.1177/2331216518769996.

27. Neff PKA, Schoisswohl S, Simoes J, Staudinger S, Langguth B, Schecklmann M, et al. Prolonged tinnitus suppression after short-term acoustic stimulation. In: Progress in Brain Research. Elsevier; 2021.Available from: https://www.sciencedirect.com/science/article/pii/S0079612321000492.

28. Roberts LE. Residual inhibition. Prog Brain Res. 2007;166:487–495. doi:10.1016/S0079-6123(07)66047-6.

29. Roberts LE, Moffat G, Baumann M, Ward LM, Bosnyak DJ. Residual Inhibition Functions Overlap Tinnitus Spectra and the Region of Auditory Threshold Shift. J Assoc Res Otolaryngol. 2008;9(4):417–435. doi:10.1007/s10162-008-0136-9.

30. Galazyuk AV, Longenecker RJ, Voytenko SV, Kristaponyte I, Nelson GL. Residual inhibition: From the putative mechanisms to potential tinnitus treatment. Hear Res. 2019;375:1–13. doi:10.1016/j.heares.2019.01.022.

31. Sedley W, Teki S, Kumar S, Barnes GR, Bamiou DE, Griffiths TD. Single-subject oscillatory gamma responses in tinnitus. Brain. 2012;135(10):3089–3100. doi:10.1093/brain/aws220.

32. Sedley W, Gander P, Kumar S, Oya H, Kovach C, Nourski K, et al. Intracranial Mapping of a Cortical Tinnitus System using Residual Inhibition. Curr Biol. 2015;25(9):1208–1214. doi:10.1016/j.cub.2015.02.075.

33. Kristeva-Feige R, Feige B, Kowalik Z, Ross B. Neuromagnetic activity during residual inhibition in tinnitus. J Audiol Med. 1995;4(3):135–142.

34. Kahlbrock N, Weisz N. Transient reduction of tinnitus intensity is marked by concomitant reductions of delta band power. BMC Biol. 2008;6(1):4. doi:10.1186/1741-7007-6-4.

35. King ROC, Singh Shekhawat G, King C, Chan E, Kobayashi K, Searchfield GD. The Effect of Auditory Residual Inhibition on Tinnitus and the Electroencephalogram. Ear and Hearing. 2021;42(1):130–141. doi:10.1097/AUD.0000000000000907.

36. Schoisswohl S, Schecklmann M, Langguth B, Schlee W, Neff P. Neurophysiological correlates of residual inhibition in tinnitus: Hints for trait-like EEG power spectra. 2021;132(7):1694–1707. doi:10.1016/j.clinph.2021.03.038.

37. Ali S, Abuhmed T, El-Sappagh S, Muhammad K, Alonso-Moral JM, Confalonieri R, et al. Explainable Artificial Intelligence (XAI): What we know and what is left to attain Trustworthy Artificial Intelligence. Information Fusion. 2023;99:101805. doi:10.1016/j.inffus.2023.101805.

38. Searchfield GD, Durai M, Linford T. A state-of-the-art review: personalization of tinnitus sound therapy. Frontiers in Psychology. 2017;8:1599.

39. Doborjeh M, Liu X, Doborjeh Z, Shen Y, Searchfield G, Sanders P, et al. Prediction of Tinnitus Treatment Outcomes Based on EEG Sensors and TFI Score Using Deep Learning. Sensors. 2023;23(2):902. doi:10.3390/s23020902.

40. Schiratti JB, Le Douget JE, Le Van Quyen M, Essid S, Gramfort A. An ensemble learning approach to detect epileptic seizures from long intracranial EEG recordings. In: 2018 IEEE International Conference on Acoustics, Speech and Signal Processing (ICASSP). IEEE; 2018. p. 856–860.

41. Craik A, He Y, Contreras-Vidal JL. Deep learning for electroencephalogram (EEG) classification tasks: a review. Journal of neural engineering. 2019;16(3):031001.

42. Donoghue T, Haller M, Peterson EJ, Varma P, Sebastian P, Gao R, et al. Parameterizing neural power spectra into periodic and aperiodic components. Nature neuroscience. 2020;23(12):1655–1665.

43. Desikan RS, Śegonne F, Fischl B, Quinn BT, Dickerson BC, Blacker D, et al. An automated labeling system for subdividing the human cerebral cortex on MRI scans into gyral based regions of interest. Neuroimage. 2006;31(3):968–980.

44. Buzsaki G. Rhythms of the Brain. Oxford university press; 2006.

45. Thomas Yeo B, Krienen FM, Sepulcre J, Sabuncu MR, Lashkari D, Hollinshead M, et al. The organization of the human cerebral cortex estimated by intrinsic functional connectivity. Journal of neurophysiology. 2011;106(3):1125–1165.

46. Li J, Zou Y, Kong X, Leng Y, Yang F, Zhou G, et al. Exploring functional connectivity alterations in sudden sensorineural hearing loss: A multilevel analysis. Brain Research. 2024;1824:148677.

47. Pedregosa F, Varoquaux G, Gramfort A, Michel V, Thirion B, Grisel O, et al. Scikit-learn: Machine learning in Python. the Journal of machine Learning research. 2011;12:2825–2830.

48. Breiman L. Random forests. Machine learning. 2001;45:5–32.

49. Friedman JH. Greedy function approximation: a gradient boosting machine. Annals of statistics. 2001; p. 1189–1232.

50. Duda RO, Hart PE, et al. Pattern classification and scene analysis. vol. 3. Wiley New York; 1973.

51. Lewis DD. Naive (Bayes) at forty: The independence assumption in information retrieval. In: European conference on machine learning. Springer; 1998. p. 4–15.

52. Hastie T, Tibshirani R, Friedman JH, Friedman JH. The elements of statistical learning: data mining, inference, and prediction. vol. 2. Springer; 2009.

53. Burges CJ. A tutorial on support vector machines for pattern recognition. Data mining and knowledge discovery. 1998;2(2):121–167.

54. Rasmussen CE, Williams CK, et al. Gaussian processes for machine learning. vol. 1. Springer; 2006.

55. Peterson LE. K-nearest neighbor. Scholarpedia. 2009;4(2):1883.

56. LeCun Y, Bottou L, Bengio Y, Haffner P. Gradient-based learning applied to document recognition. Proceedings of the IEEE. 1998;86(11):2278–2324.

57. Boser BE, Guyon IM, Vapnik VN. A training algorithm for optimal margin classifiers. In: Proceedings of the fifth annual workshop on Computational learning theory; 1992. p. 144–152.

58. Lundberg SM, Lee SI. A Unified Approach to Interpreting Model Predictions. In: Guyon I, Luxburg UV, Bengio S, Wallach H, Fergus R, Vishwanathan S, et al., editors. Advances in Neural Information Processing Systems 30. Curran Associates, Inc.; 2017. p. 4765–4774. Available from: http://papers.nips.cc/paper/7062-a-unified-approach-to-interpreting-model-predictions.pdf.

59. Jolliffe IT. Principal component analysis for special types of data. Springer; 2002.

60. Weinberger KQ, Saul LK. Distance metric learning for large margin nearest neighbor classification. Journal of machine learning research. 2009;10(2).

61. Gini CW. Variability and mutability, contribution to the study of statistical distributions and relations. Studi Economico-Giuridici della R Universita de Cagliari. 1912;.

62. Adjamian P, Sereda M, Zobay O, Hall DA, Palmer AR. Neuromagnetic Indicators of Tinnitus and Tinnitus Masking in Patients with and without Hearing Loss. J Assoc Res Otolaryngol. 2012;13(5):715–731. doi:10.1007/s10162-012-0340-5.

63. Lorenz I, Müller N, Schlee W, Hartmann T, Weisz N. Loss of alpha power is related to increased gamma synchronization-A marker of reduced inhibition in tinnitus? Neurosci Lett. 2009;453(3):225–228. doi:10.1016/j.neulet.2009.02.028.

64. Balkenhol T, Wallhäusser-Franke E, Delb W. Psychoacoustic Tinnitus Loudness and Tinnitus-Related Distress Show Different Associations with Oscillatory Brain Activity. PLoS One. 2013;8(1):e53180. doi:10.1371/journal.pone.0053180.

65. van der Loo E, Gais S, Congedo M, Vanneste S, Plazier M, Menovsky T, et al. Tinnitus intensity dependent gamma oscillations of the contralateral auditory cortex. PLoS One. 2009;4(10):e7396. doi:10.1371/journal.pone.0007396.

66. Sedley W, Cunningham MO. Do cortical gamma oscillations promote or suppress perception? An under-asked question with an over-assumed answer. Frontiers in human neuroscience. 2013;7:595. doi:10.3389/fnhum.2013.00595.

67. Uhlhaas PJ, Singer W. Neural Synchrony in Brain Disorders: Relevance for Cognitive Dysfunctions and Pathophysiology. Neuron. 2006;52(1):155–168. doi:10.1016/j.neuron.2006.09.020.

68. Ray S, Niebur E, Hsiao SS, Sinai A, Crone NE. High-frequency gamma activity (80–150 Hz) is increased in human cortex during selective attention. Clinical Neurophysiology. 2008;119(1):116 – 133. doi:10.1016/j.clinph.2007.09.136.

69. Schnitzler A, Gross J. Normal and pathological oscillatory communication in the brain. Nature Reviews Neuroscience. 2005;6(4):285–296. doi:10.1038/nrn1650.

70. Sedley W, Friston KJ, Gander PE, Kumar S, Griffiths TD. An Integrative Tinnitus Model Based on Sensory Precision. Trends in Neurosciences. 2016;39(12):799 – 812. doi:10.1016/j.tins.2016.10.004.

71. Engel AK, Fries P, Singer W. Dynamic predictions: Oscillations and synchrony in top–down processing. Nature Reviews Neuroscience. 2001;2(10):704–716. doi:10.1038/35094565.

72. Schlee W, Schecklmann M, Lehner A, Kreuzer PM, Vielsmeier V, Poeppl TB, et al. Reduced Variability of Auditory Alpha Activity in Chronic Tinnitus. Neural Plast. 2014;2014:1–9. doi:10.1155/2014/436146.

73. Sedley W, Parikh J, Edden RAE, Tait V, Blamire A, Griffiths TD. Human Auditory Cortex Neurochemistry Reflects the Presence and Severity of Tinnitus. The Journal of neuroscience : the official journal of the Society for Neuroscience. 2015;35(44):14822 – 14828. doi:10.1523/jneurosci.2695-15.2015.

74. Isler B, Burg Nv, Kleinjung T, Meyer M, Stämpfli P, Zölch N, et al. Lower glutamate and GABA levels in auditory cortex of tinnitus patients: a 2D-JPRESS MR spectroscopy study. Scientific Reports. 2022;12(1):4068. doi:10.1038/s41598-022-07835-8.

75. Lange J, Keil J, Schnitzler A, Dijk Hv, Weisz N. The role of alpha oscillations for illusory perception. Behavioural brain research. 2014;271:294 – 301. doi:10.1016/j.bbr.2014.06.015.

76. Llińas RR, Ribary U, Jeanmonod D, Kronberg E, Mitra PP. Thalamocortical dysrhythmia: A neurological and neuropsychiatric syndrome characterized by magnetoencephalography. PNAS USA. 1999;96(26):15222–15227. doi:10.1073/pnas.96.26.15222.

77. Llińas R, Urbano FJ, Leznik E, Ramírez RR, van Marle HJF. Rhythmic and dysrhythmic thalamocortical dynamics: GABA systems and the edge effect. Trends Neurosci. 2005;28(6):325–333. doi:10.1016/j.tins.2005.04.006.

78. De Ridder D, Vanneste S, Langguth B, Llinas R. Thalamocortical Dysrhythmia: A Theoretical Update in Tinnitus. Front Neurol. 2015;6. doi:10.3389/fneur.2015.00124.

79. Zatorre RJ, Belin P. Spectral and Temporal Processing in Human Auditory Cortex. Cerebral Cortex. 2001;11(10):946 – 953. doi:10.1093/cercor/11.10.946.

80. Jensen O, Mazaheri A. Shaping Functional Architecture by Oscillatory Alpha Activity: Gating by Inhibition. Frontiers in human neuroscience. 2010;4. doi:10.3389/fnhum.2010.00186.

81. Rauschecker JP, Leaver AM, Mühlau M. Tuning Out the Noise: Limbic-Auditory Interactions in Tinnitus. Neuron. 2010;66(6):819 – 826. doi:10.1016/j.neuron.2010.04.032.

82. Song JJ, Vanneste S, Ridder DD. Dysfunctional noise cancelling of the rostral anterior cingulate cortex in tinnitus patients. PLoS ONE. 2015;10(4):e0123538. doi:10.1371/journal.pone.0123538.

83. Meyer M, Neff P, Liem F, Kleinjung T, Weidt S, Langguth B, et al. Differential tinnitus-related neuroplastic alterations of cortical thickness and surface area. Hearing Research. 2016;342:1 – 12. doi:10.1016/j.heares.2016.08.016.

84. Rauschecker JP, May ES, Maudoux A, Ploner M. Frontostriatal Gating of Tinnitus and Chronic Pain. Trends in cognitive sciences. 2015;19(10):567 – 578. doi:10.1016/j.tics.2015.08.002.

85. Berger JI, Billig AJ, Sedley W, Kumar S, Griffiths TD, Gander PE. What is the role of the hippocampus and parahippocampal gyrus in the persistence of tinnitus? Human Brain Mapping. 2024;45(3):e26627. doi:10.1002/hbm.26627.

86. Lee SY, Chang M, Kwon B, Choi BY, Koo JW, Moon T, et al. Is the posterior cingulate cortex an on-off switch for tinnitus?: A comparison between hearing loss subjects with and without tinnitus. Hearing Research. 2021;411:108356. doi:10.1016/j.heares.2021.108356.

87. Knight RT, Scabini D, Woods DL. Prefrontal cortex gating of auditory transmission in humans. Brain Research. 1989;504(2):338–342. doi:10.1016/0006-8993(89)91381-4.

88. Vanneste S, Plazier M, Loo Evd, Heyning PVd, Ridder DD. The differences in brain activity between narrow band noise and pure tone tinnitus. PLoS ONE. 2010;5(10):e13618.

89. Vanneste S, Plazier M, Loo Evd, Heyning PVd, Ridder DD. The difference between uni- and bilateral auditory phantom percept. Clinical Neurophysiology. 2011;122(3):578 – 587. doi:10.1016/j.clinph.2010.07.022.

90. Lee SJ, Park J, Lee SY, Koo JW, Vanneste S, Ridder DD, et al. Triple network activation causes tinnitus in patients with sudden sensorineural hearing loss: A model-based volume-entropy analysis. Frontiers in Neuroscience. 2022;16:1028776. doi:10.3389/fnins.2022.1028776.

91. Schlee W, Hartmann T, Langguth B, Weisz N. Abnormal resting-state cortical coupling in chronic tinnitus. BMC Neurosci. 2009;10(1):11. doi:10.1186/1471-2202-10-11.

92. Schaette R, Kempter R. Computational models of neurophysiological correlates of tinnitus. Frontiers in Systems Neuroscience. 2012;6:34. doi:10.3389/fnsys.2012.00034.

93. Roberts LE, Eggermont JJ, Caspary DM, Shore SE, Melcher JR, Kaltenbach JA. Ringing ears: the neuroscience of tinnitus. The Journal of neuroscience : the official journal of the Society for Neuroscience. 2010;30(45):14972 – 14979. doi:10.1523/jneurosci.4028-10.2010.

94. Tzounopoulos T, Balaban C, Zitelli L, Palmer C. Towards a Mechanistic-Driven Precision Medicine Approach for Tinnitus. Journal of the Association for Research in Otolaryngology: JARO. 2019;20(2):115–131. doi:10.1007/s10162-018-00709-9.

95. Jackson R, Vijendren A, Phillips J. Objective Measures of Tinnitus: a Systematic Review. Otology & Neurotology. 2019;40(2):154–163. doi:10.1097/mao.0000000000002116.

96. Edvall NK, Mehraei G, Claeson M, Lazar A, Bulla J, Leineweber C, et al. Alterations in auditory brainstem response distinguish occasional and constant tinnitus. Journal of Clinical Investigation. 2022;doi:10.1172/jci155094.

